# Shaping and interpretation of Dpp morphogen gradient by endocytic trafficking

**DOI:** 10.1101/2023.03.27.534445

**Authors:** Sheida Hadji Rasouliha, Gustavo Aguilar, Cindy Reinger, Shinya Matsuda

**Affiliations:** Growth & Development, Biozentrum, Spitalstrasse 41, University of Basel, 4056 Basel, Switzerland

## Abstract

Dpp/BMP is a morphogen that controls patterning and growth in the Drosophila wing disc. Endocytic trafficking has been proposed to control extracellular Dpp spreading or removal, and Dpp signaling. However, how Dpp morphogen gradient is shaped and interpreted by endocytic trafficking remains unclear due to the lack of tools to visualize Dpp distribution at the physiological level. Here we generate functional fluorescent protein tagged *dpp* alleles to visualize both extracellular and intracellular Dpp distribution. Using these alleles, we found that, while Dynamin-mediated internalization of Dpp is required for Dpp signal activation, Rab5-mediated trafficking is not required for Dpp spreading or signaling as proposed before but for shutting off Dpp signaling through downregulating activated receptors. We provide evidence that Dpp signaling is terminated at Multivesicular body (MVB) likely through sorting activated receptors into intraluminal vesicles (ILVs) rather than Rab7-mediated lysosomal degradation. We further found that blocking MVB formation expanded Dpp signaling gradient without affecting extracellular Dpp gradient, thus compromising extracellular Dpp gradient interpretation. These results indicate that extracellular Dpp gradient is shaped by Dynamin-mediated internalization of Dpp and interpreted by duration of Dpp signaling.

## Introduction

Morphogens are signaling molecules that are produced by a localized source of cells, and control the fate of their neighboring cells in a concentration dependent manner ^1^. Among morphogens, Decapentaplegic (Dpp), the homologue of the vertebrate bone morphogenetic protein 2/4 (BMP2/4) has served as an excellent model system to understand morphogen function.

Dpp is produced in a stripe of cells in the anterior compartment along the anterior/posterior compartment boundary of the wing imaginal disc and controls patterning and growth of the Drosophila wing. From the source cells, Dpp spreads and forms a concentration gradient in the tissue ^2–6^. Given the severe patterning and growth defects in *dpp* mutant flies, Dpp spreading from the stripe of cells has been thought to be essential for patterning and growth. However, it has recently been shown that blocking Dpp spreading from the source cells in the wing discs severely affected the posterior patterning and growth without greatly affecting anterior patterning and growth ^7^, indicating that Dpp spreading is not as important as previously expected. Nevertheless, a variety of extracellular and cell surface molecules have been shown to play essential roles in Dpp gradient formation and signaling gradient.

Dpp is thought to bind to the Type I and Type II receptors, Tkv and Punt, on the cell surface to induce phosphorylation of Mad (pMad) in the target tissue. pMad is then translocated into the nucleus to control the expression of Dpp target genes mainly by repressing Brk, which acts as a repressor for Dpp target genes ^4^. Thus, the graded extracellular Dpp gradient is converted into the nuclear pMad gradient and the inverse-in-shape Brk gradient, which regulates the nested expression of the target genes to specify the position of the future adult wing veins such as L2 and L5 as well as growth ^8–10^.

In addition to extracellular regulation and nuclear interpretation, endocytic trafficking has been implicated in shaping and interpretation of gradients of different morphogens ^11–14^. However, how the extracellular Dpp morphogen gradient is shaped and interpreted by endocytic trafficking remains unclear.

Several models have been proposed for the role of endocytosis on Dpp morphogen gradient. First, since Dpp accumulated in *tkv* mutant clones especially in cells close to the source cells, Dpp was thought to be internalized and transported by Tkv through repeated cycles of endocytosis and exocytosis ^15^. Second, it has recently been proposed that heparan sulfate proteoglycans such as Dally, but not Tkv, acts as a cell-surface receptor to internalize and recycle Dpp to contribute to the extracellular Dpp morphogen gradient ^16^. In this case, Dpp is thought to bind to Tkv intracellularly to activate Dpp signaling. Although both models have been challenged ^17–20^, endocytic trafficking may control Dpp spreading through other cell surface factors. Third, Tkv-mediated endocytosis has been proposed to simply acts as a sink to remove extracellular Dpp ^20–22^, and Dally antagonizes this process to establish a long range Dpp gradient ^20,21^. Thus, regardless of the role of extracellular and cell-surface factors on regulation of the extracellular Dpp gradient, if and how endocytic trafficking itself influences the extracellular Dpp gradient remains controversial. In addition, how extracellular Dpp gradient is interpreted at the cellular level remains unknown. Interestingly, it has been shown that Dpp mainly exists intracellularly and Dpp signaling is lost in endocytosis deficient cells ^23–26^, indicating the importance of internalized Dpp for its signal activation. However, it remains unclear in which endocytic compartment Dpp signaling is activated and shut off, and whether the duration of Dpp signaling is required to interpret the extracellular Dpp gradient. Recently, fluorophore-conjugated anti-GFP nanobody was used to label and trace only the internalized GFP-Dpp ^16^. However, it remains unclear if nanobody-bound GFP-Dpp is functional. Thus, the role of endocytosis on Dpp morphogen gradient formation and interpretation remains unclear partly due to the lack of *dpp* alleles to visualize both extracellular and intracellular Dpp distribution at the physiological level.

In this study, we generated such functional fluorescent protein tagged *dpp* alleles and systematically investigated the role of endocytic trafficking on Dpp morphogen gradient formation and Dpp signaling activity. We found that while blocking Dynamin-mediated endocytosis expands Dpp distribution and impairs Dpp signaling, blocking Rab5-mediated early endosomal formation expanded Dpp signaling range due to impaired downregulation of Tkv. This indicated that Dpp signaling is activated upon Dynamin-mediated internalization of Dpp and terminated through Rab5-mediated endocytic trafficking. We showed that blocking the multivesicular body (MVB) formation, but not Rab7-mediated lysosomal degradation, expanded intracellular Dpp distribution and Dpp signaling range, but extracellular Dpp gradient remained unaffected. These results indicate that termination of Dpp signaling at the MVB converts extracellular Dpp gradient into Dpp signaling gradient. Taken together, these results suggest that the extracellular Dpp gradient is shaped by Dynamin-mediated internalization of Dpp and interpreted by duration of Dpp signaling.

## Results

### Visualization of extracellular and intracellular Dpp gradient in the wing disc

We previously generated an endogenous GFP-dpp allele by inserting GFP after the last processing sites of Dpp to tag all the mature Dpp ^7^. However, the resulting GFP-Dpp signal was too weak to visualize the graded distribution (Fig. 1A). Similar results were obtained using the other GFP-dpp allele ^16^. To better visualize the endogenous Dpp gradient, we then inserted mGreenLantern (mGL) ^27^ or mScarlet (mSC) ^28^ into the *dpp* locus to generate endogenous *mGL-dpp* and *mSC-dpp* alleles respectively. Interestingly, we found that the mGL-Dpp shows brighter fluorescent signal than GFP-Dpp (Fig. 1A, B) and a graded distribution outside the stripe of Dpp source cells (Fig. 1B). Similar graded distribution of mSC-dpp signal was also observed (Fig. 1C).

**Figure 1:**
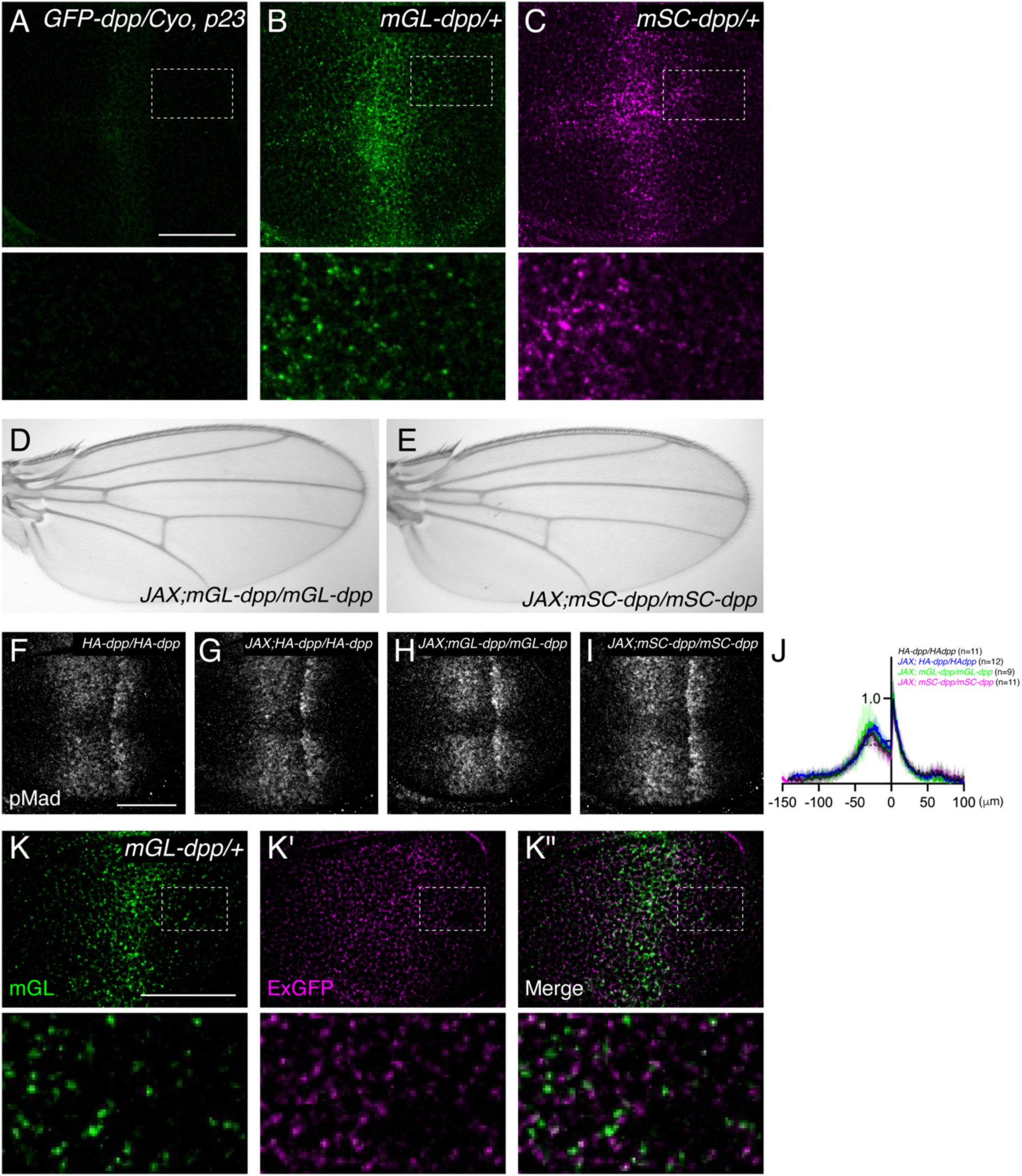
Visualization of endogenous Dpp morphogen gradient in the wing disc. (A) GFP-Dpp signal from *GFP-dpp/Cyo, p23*. (A), mGL-Dpp signal from *mGL-dpp/+* wing disc mSC-Dpp signal from *mSC-dpp/+*. (D) Adult wing of *JAX; mGL-dpp/mGL-dpp*. (E) Adult wing of JAX; *mSC-dpp/mSC-dpp*. (F-I) α-pMad staining of *HA-dpp/HA-dpp* (F), *JAX; HA-dpp/HA-dpp* (G), *JAX; mGL-dpp/mGL-dpp* (H), *JAX; mSC-dpp/mSC-dpp* (I) wing disc. (J) Average fluorescence intensity profile of (F-I). Data are presented as mean +/-SD. (K) mGL-Dpp signal (K), extracellular a-GFP staining (K’), and merge (K”) of *mGL-dpp/+* wing disc. Scale Bar: 50um.

Unlike the *GFP-dpp* allele, the two newly generated alleles were not haploinsufficient but semi-lethal. To overcome the partial embryonic lethality, we introduced a transgene called “JAX”, which contains the genomic region of *dpp* critical for the early embryogenesis ^29^ but does not rescue the wing phenotypes of *dpp* mutants ^20^. We found that the lethality of each allele was greatly rescued by JAX and became easily homozygous viable without obvious phenotypes (Fig. 1D, E). JAX did not affect Dpp signal in the functional *HA-dpp* ^7^ wing discs (Fig. 1F, G, J) and Dpp signal was comparable between *JAX;HA-dpp*, *JAX;mGL-dpp* and *JAX;mSC-dpp* wing discs (Fig. 1G-J). These results suggest that *mGL-dpp* and *mSC-dpp* allele are functional at least during wing disc development.

To address whether the mGL-Dpp signal is derived from extracellular or intracellular Dpp, we performed extracellular staining using anti-GFP antibody and compared the extracellular mGL-Dpp distribution with the total mGL-Dpp fluorescent signal. In contrast to the higher total mGL-Dpp signal in the center of wing disc, where *dpp* is expressed (Fig. 1K), the extracellular mGL-Dpp showed a shallow graded distribution (Fig. 1K’). The two signals rarely colocalize (Fig. 1K”), indicating that the majority of mGL-Dpp signal is derived from the intracellular Dpp. Consistently, the extracellular mGL-Dpp distribution, but not the total mGL-Dpp fluorescent signal, was sensitive to the acid wash, which efficiently removes extracellular proteins ^16^ (Fig. S1).

To test where the endogenous Dpp is localized within the cells, we compared the endogenous mSC-Dpp localization with different Rab proteins tagged with eYFP (Fig. 2). The Mander’s coefficient (M1) revealed that mSC-Dpp colocalizes with the early endosome marker Rab5-eYFP (Fig. 2A’), the late endosome marker Rab7-eYFP (Fig. 2B’), the fast-recycling endosome marker Rab4-eYFP (Fig. 2C’), and the slow recycling endosome marker Rab11-eYFP (Fig. 2D’) to different extents, showing that the internalized Dpp is trafficked to different endocytic compartments.

**Figure 2:**
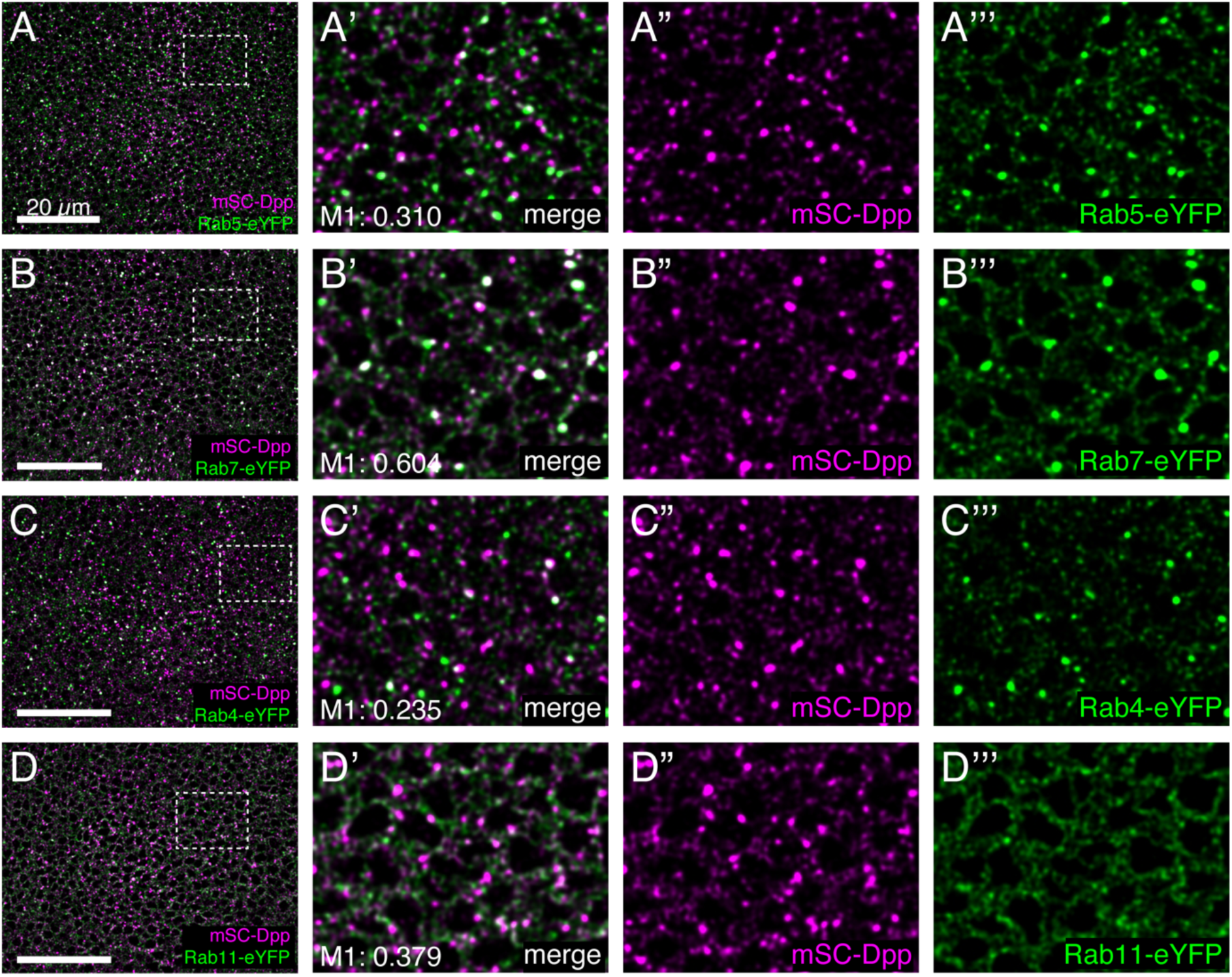
Colocalization of mSC-Dpp with different Rabs. (A-D) Comparison of mSC-Dpp signal with Rab5-eYFP (A), Rab7-eYFP (B), Rab4-eYFP (C), Rab11-eYFP (D) in the late third instar wing imaginal discs. Mander’s coefficient (M1) indicates the percentage of overlap of mSC-Dpp with different Rabs.

### Rab5 is required for downregulating Dpp signal

To study how different endocytic compartments contribute to Dpp gradient formation and signaling, we first knocked down Dynamin GTPase (*Drosophila* homologue: shibire), a critical factor to excise the formed vesicles and separate them from the plasma membrane ^30^ Consistent with the idea that Dpp signaling is activated upon endocytosis ^22^, we found that the temperature-sensitive allele of shibire (*shi^ts^*^1^) led to a complete loss of Dpp signaling at restrictive temperatures for 2h (Fig. 3A-C). Previous studies showed that loss of Rab5 by dominant negative form of Rab5 also reduces Dpp signaling and its target gene expression, indicating that Dpp is transported through endocytosis ^15^, and/or Dpp signaling is activated at the level or downstream of the early endosome ^31^. In stark contrast, we found that temporal knocking down of Rab5 by RNAi using the temperature-sensitive Gal80 (tubGal80ts) in the dorsal compartment of the wing discs resulted in an increase in Dpp signaling activity compared with the control ventral compartment (Fig. 3D-F). Similar results were obtained using different RNAi lines against Rab5 or the dominant negative form of Rab5 (Fig.3, G-J). Inducing *rab5* null clones (*rab5*^2^)^32^ also led to a cell autonomous reduction in Brk intensity (Fig. 3K-K’), consistent with an increase of Dpp signaling. These results suggest that Dpp signaling activated upon endocytosis is downregulated through Rab5-mediated trafficking.

**Figure 3:**
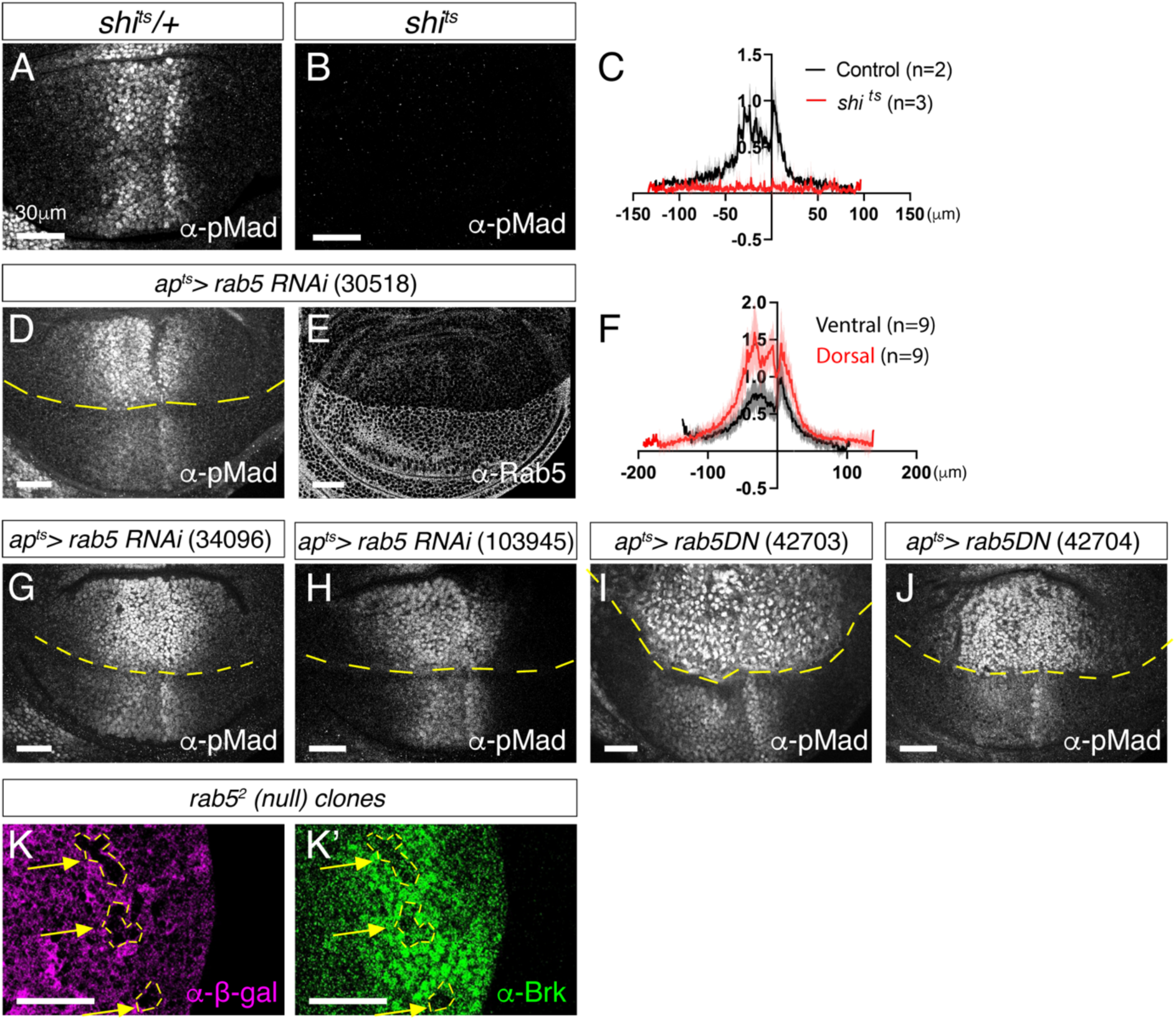
Rab5 is required for downregulating Dpp signaling. (A-B) α-pMad staining of *shi^ts^/+* wing disc (control) (A) and *shi^ts^* wing disc (B) upon 2h at restrictive temperatures. (C) Average fluorescence intensity profile of (A, B). Data are presented as mean +/-SD. (D, E) α-pMad staining (D) and α-Rab5 staining (E) of *ap^ts^>rab5RNAi* (30518). (F) Average fluorescence intensity profile of (D). Data are presented as mean +/-SD. (G-J) α-pMad staining of *ap^ts^>rab5RNAi* (34096) (G), *ap^ts^>rab5RNAi* (103945) (H), *ap^ts^>rab5DN* (42703) (I), and *ap^ts^>rab5DN* (42704) (J). (K) *rab5*^2^ null clones generated in the peripheral regions visualized via absence of α-β-gal staining (K) and a-Brk staining (K’). Scale bar:30μm.

### The effects of Rab5 on Dpp distribution

How does loss of Rab5 lead to an increase in Dpp signaling? To test if *dpp* is involved, we knocked down Rab5 in the dorsal compartment in *dpp, brk* double mutants, in which the wing disc could grow in the absence of *dpp*. We found that pMad was not upregulated under this condition, indicating that the observed phenotype was dependent on Dpp (Fig. 4A). To test if changes in *dpp* transcription were involved, we then followed a *dpp* transcription reporter *dpp-lacZ* upon knocking down Rab5 via RNAi and found no changes in *dpp* transcription (Fig. 4B), indicating that upregulation of *dpp* transcription was not the cause of increase of Dpp signaling upon knocking down Rab5.

**Figure 4:**
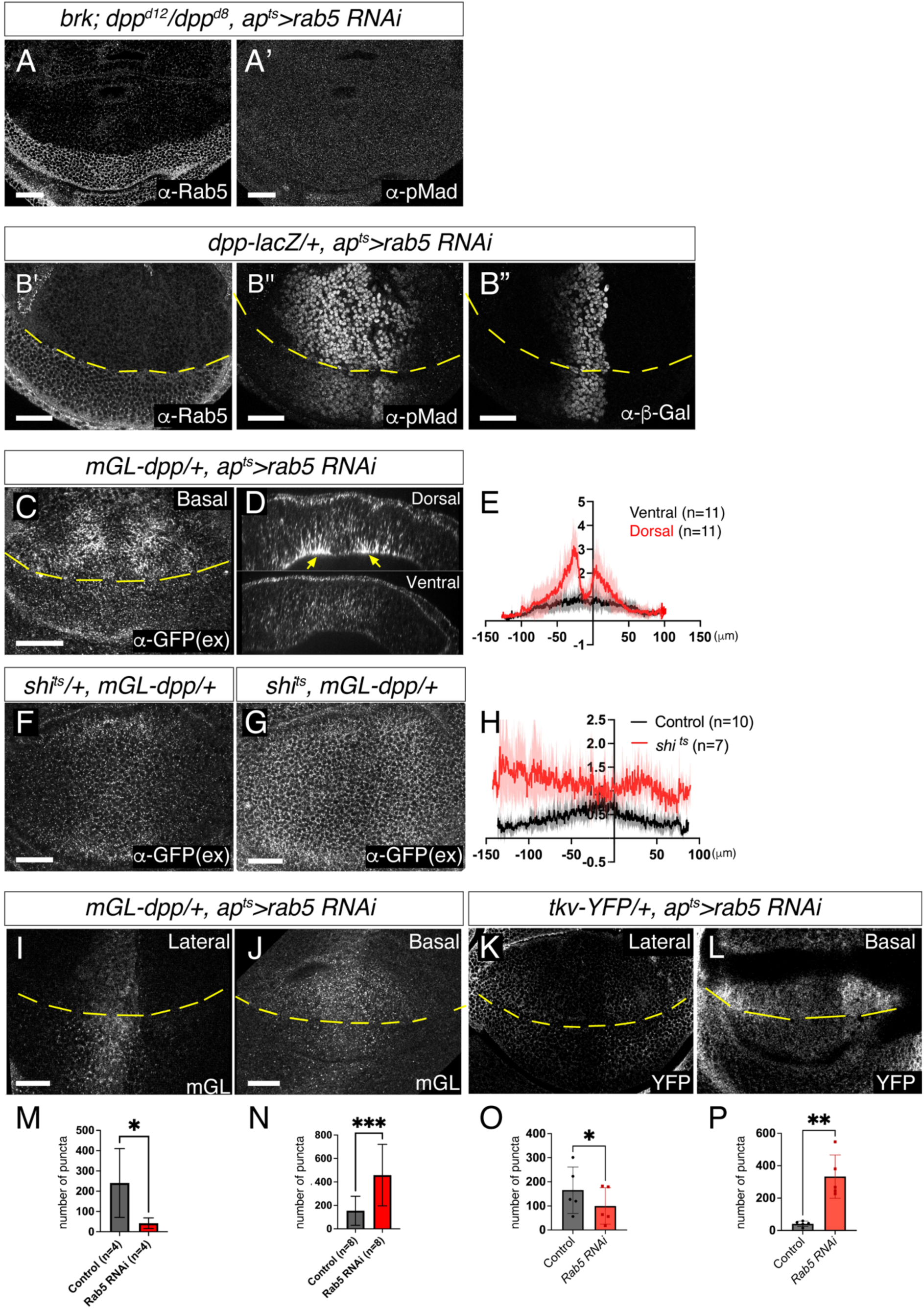
Changes in Dpp distribution in absence of Rab5. (A) α-Rab5 staining (A) and α-pMad staining (A’) in *brk, dpp^d^*^8^*/dpp^d^*^12^*, ap^ts^>Rab5 RNAi* wing disc. (B) a-Rab5 staining (B), α-pMad staining (B’), and α-β-gal staining (B”) in *dpp-lacZ/+, ap^ts^>Rab5 RNAi*. (C) Extracellular α-GFP staining in *mGL-dpp/+, ap^ts^>Rab5 RNAi*. (D) Optical cross-section of (C). (E) Average fluorescence intensity profile of (C). Data are presented as mean +/-SD. (F, G) Extracellular α-GFP staining in *shi^ts^/+, mGL-dpp/+* wing disc (F) and *shi^ts^, mGL-dpp/+* (G) after 2h at restrictive temperature of 34°C. (H) Average fluorescence intensity profile of (F, G). Data are presented as mean +/-SD. (I, J) mGL-Dpp intracellular signal in the lateral side (I) and basal side (J) in *mGL-dpp/+, ap^ts^>rab5 RNAi* wing disc. (K, L) Tkv-YFP (total) signal of lateral side (K) and basal side (L) of *tkv-YFP/+, ap^ts^>rab5 RNAi* wing disc. (M-P) Comparison of the number of puncta of (I-L). Ratio-paired t-test with p<0.05 was used for the comparison; p=0.0383 (n=4) (M), p=0.0001 (n=8) (N), p=0.0123 (n=5) (O) p=0.0010 (n=5) (P). Scale bar: 30μm

We then asked if the changes in Dpp distribution and/or trafficking affect Dpp signaling in this condition. Consistent with impaired endocytosis in *rab5* mutants ^33^, loss of Rab5 resulted in increase of the extracellular mGL-Dpp (Fig. 4, C-E) similar to *shi^ts^*^1^ (Fig. 4F-H). Visualizing cross-sections from the wing imaginal discs showed that the extracellular mGL-Dpp increased in the basolateral side, especially outside of the Dpp producing cells (Fig. 4D, yellow arrowheads). This increase of extracellular Dpp could lead to activation of Dpp signaling. However, loss of Dpp signaling in *shi^t^*^s1^ despite accumulation of extracellular mGL-Dpp (Fig. 3B, 4F-H) suggests that Dynamin-mediated endocytosis of Dpp is required for Dpp signal activation. Thus, the accumulation of the extracellular Dpp by loss of Rab5 is unlikely the direct cause of increase in Dpp signaling activity.

Although impaired, internalization has been shown to occur and the following endosomal maturation is blocked in Rab5 mutants ^32–34^. Consistently, we found that loss of Rab5 by RNAi decreased intracellular mGL-Dpp puncta in the lateral side (Fig. 4I, M) but increased in more basal side of the disc (Fig. 4J, N). Similarly, Tkv-YFP puncta decreased in the lateral side (Fig. 4K, O), but increased in the basal side of the wing discs in absence of Rab5 (Fig. 4L, P). These results raise a possibility that Dpp signal is activated upon internalization of Dpp but not terminated in the absence of Rab5, thereby expanding Dpp signaling.

### Rab5-mediaed trafficking terminates Dpp signaling through downregulating Tkv

If the activated Tkv signal is not terminated in the absence of Rab5, the increased pMad intensity by loss of Rab5 should be rescued by removal of Tkv. To test this, we applied deGradHA, a genetically encoded method to artificially degrade HA-tagged proteins ^35^. Since Tkv is the critical receptor for Dpp signaling, we used the deGradHA tool to degrade only one copy of Tkv-HA-eGFP in the dorsal compartment of the wing discs (Fig. 5). While pMad intensity was similar between the dorsal and the ventral compartment in the control wing discs (Fig. 5A), knocking down Rab5 via RNAi in the dorsal compartment led to an increase in pMad intensity compared to the ventral compartment (Fig. 5B). In contrast, simultaneous knocking down of Rab5 via RNAi and degrading one copy of Tkv-HA-eGFP via deGradHA in the dorsal compartment rescued the dorsal pMad intensity comparable to the ventral pMad signal (Fig. 5C). Partial degradation of one copy of Tkv-HA-eGFP via deGradHA alone did not affect the pMad gradient except a slight decrease along the A/P compartment boundary, likely due to the low levels of Tkv in that region (Fig. 5D). These results suggest that Dpp signaling is not terminated in the absence of Rab5 due to impaired downregulation of activated Tkv.

**Figure 5:**
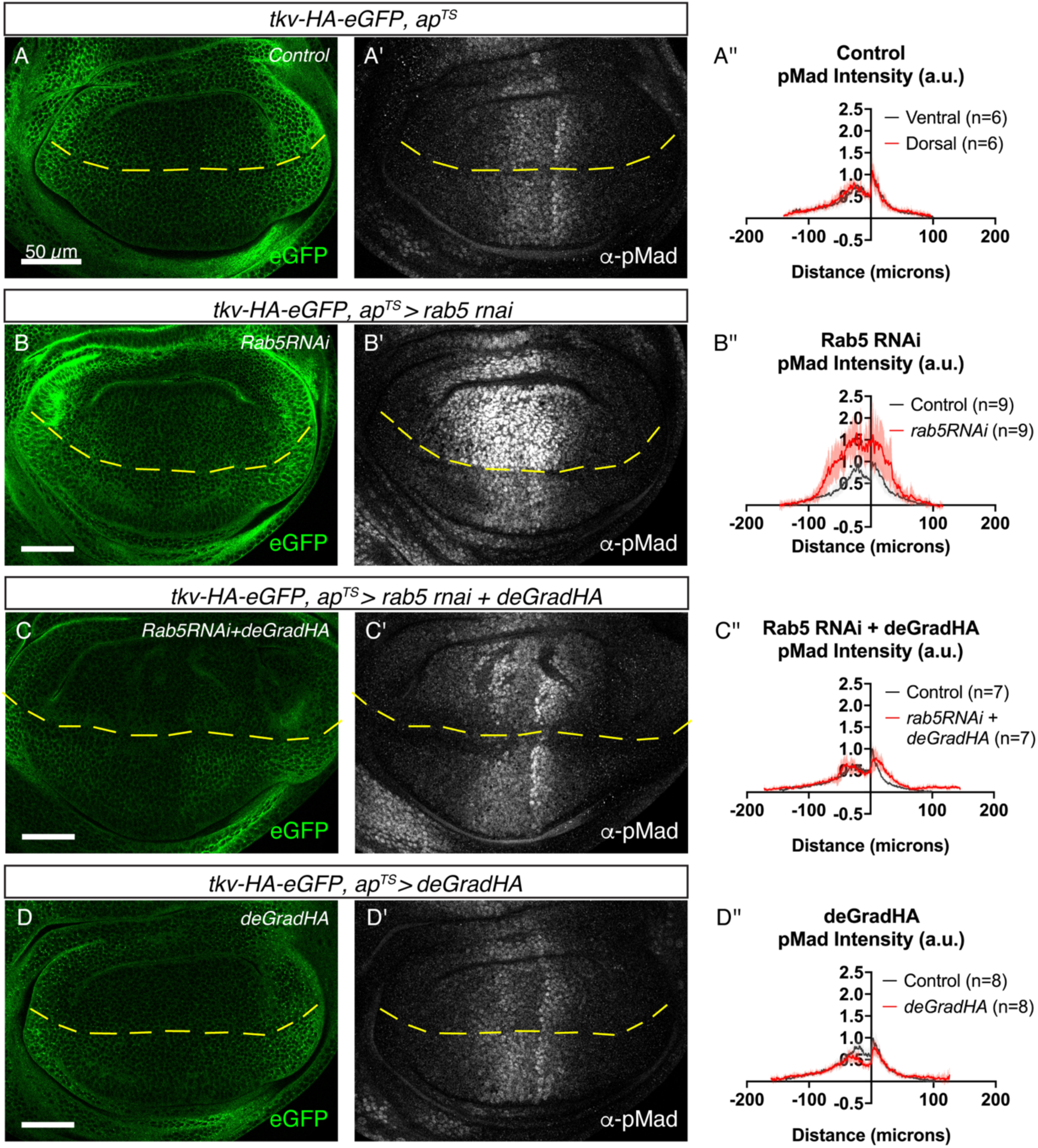
Partial degradation of Tkv rescues increase of Dpp signaling upon loss of Rab5. (A-D) Tkv-HA-eGFP signal (A-D) and α-pMad staining (A’-D’) of *tkv-HA-eGFP/+, ap^ts^>+* wing disc (Control) (A), *tkv-HA-eGFP/+, ap^ts^>rab5 RNAi* wing disc (B), *tkv-HA-eGFP/+, ap^ts^>rab5 RNAi, deGradHA* wing disc (C) and *tkv-HA-eGFP/+, ap^ts^>deGradHA* wing disc (D). (A”-D”) Average fluorescence intensity profiles of (A’-D’). Data are presented as mean +/-SD. Scale bar: 50μm

### MVBs formation terminates Dpp signaling

We then asked in which endocytic trafficking Dpp signal is terminated? As the early endosome matures to the late endosome, the ESCRT components recognize and sort ubiquitinated proteins into the intraluminal vesicles (ILVs) to form the multivesicular bodies (MVBs). The late endosome containing MVBs are then fused with lysosome to degrade the contents of MVBs^36^. It has been proposed that Dpp signaling is shut off through endosomal degradation of activated Tkv ^37,38^. Indeed, we found that knocking down factors required for MVB formation such as ESCRT-II component TSG101, ESCRT-III component Shrub, or Vps4 by RNAi in the dorsal compartment led to an increase in range and intensity of the pMad signal compared to the ventral compartment (Fig. 6A-C). Consistent with the defects in sorting of ubiquitinated receptors into ILVs, Tkv and ubiquitin accumulated and highly colocalized upon knocking down Vps4, Shrub, or Tsg101 (Fig. 6D, Fig. S2), but not upon knocking down Rab7 (Fig. 6E). We then tested if blocking formation of MVBs affects extracellular and/or intracellular Dpp distribution. Consistent with the defects in sorting of Dpp, we found that the intracellular mGL-Dpp accumulated as large puncta, but extracellular mGL-Dpp gradient remained unaffected upon knocking down Vps4 (Fig. 6F-H). These results suggest that termination of Dpp signaling at the level or downstream of MVB formation is involved in converting the extracellular Dpp gradient into the Dpp signaling gradient.

**Figure 6.**
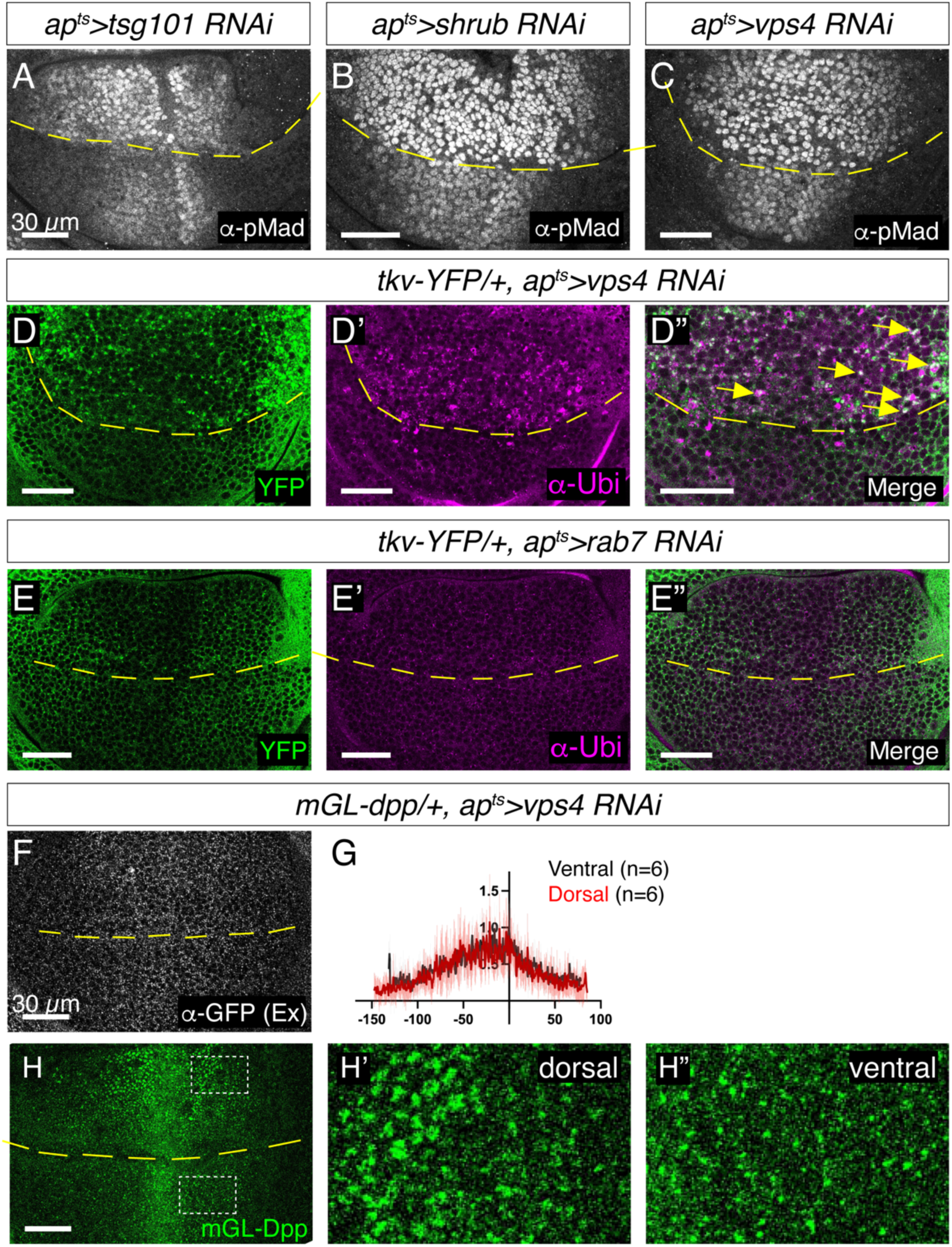
Formation of MVBs is required for Dpp signaling termination. (A-C) α-pMad staining of *ap^ts^>tsg101 RNAi* (A), *ap^ts^>shrub RNAi* (B), and *ap^ts^>Vps4 RNAi* (C) wing disc. (D) Tkv-YFP signal (D), α-Ubiquitin staining (D’), and merge (D”) in *tkv-YFP/+, ap^ts^>vps4 RNAi* wing disc. (E) Tkv-YFP signal (E), α-Ubiquitin staining (E’), and merge (E”) in *tkv-YFP/+, ap^ts^>rab7 RNAi* wing disc. (F) Extracellular a-GFP staining of *mGL-dpp/+, ap^ts^>Vps4 RNAi* wing disc. (G) Average fluorescence intensity profiles of (F). Data are presented as mean +/-SD. (H) mGL-Dpp signal from apical side (H), with magnified region in the dorsal compartment (H’), and the ventral compartment (H”) of *mGL-dpp/+, ap^ts^>Vps4 RNAi* wing disc. (H’’’) Comparison of size of mGL-Dpp puncta in H’ and H’’, ratio-paired t-test with p<0.05 was used for the comparison; p=0.0325 (n=5).

### Late endosomal trafficking is dispensable for terminating Dpp signaling

We then addressed if Dpp signaling is terminated through lysosomal degradation. Surprisingly, inducing clones of cells mutant for *rab7* (null mutant) ^39^ or knocking down Rab7 by RNAi (Fig. 7A-D) reduced Rab7 but did not affect Dpp signaling activity. The extracellular and the intracellular mGL-Dpp signal also remained unchanged upon knocking down Rab7 by RNAi (Fig. 7E-G). These results suggest that MVB formation, but not Rab7-mediated lysosomal degradation, is critical to terminate Dpp signaling.

**Figure 7:**
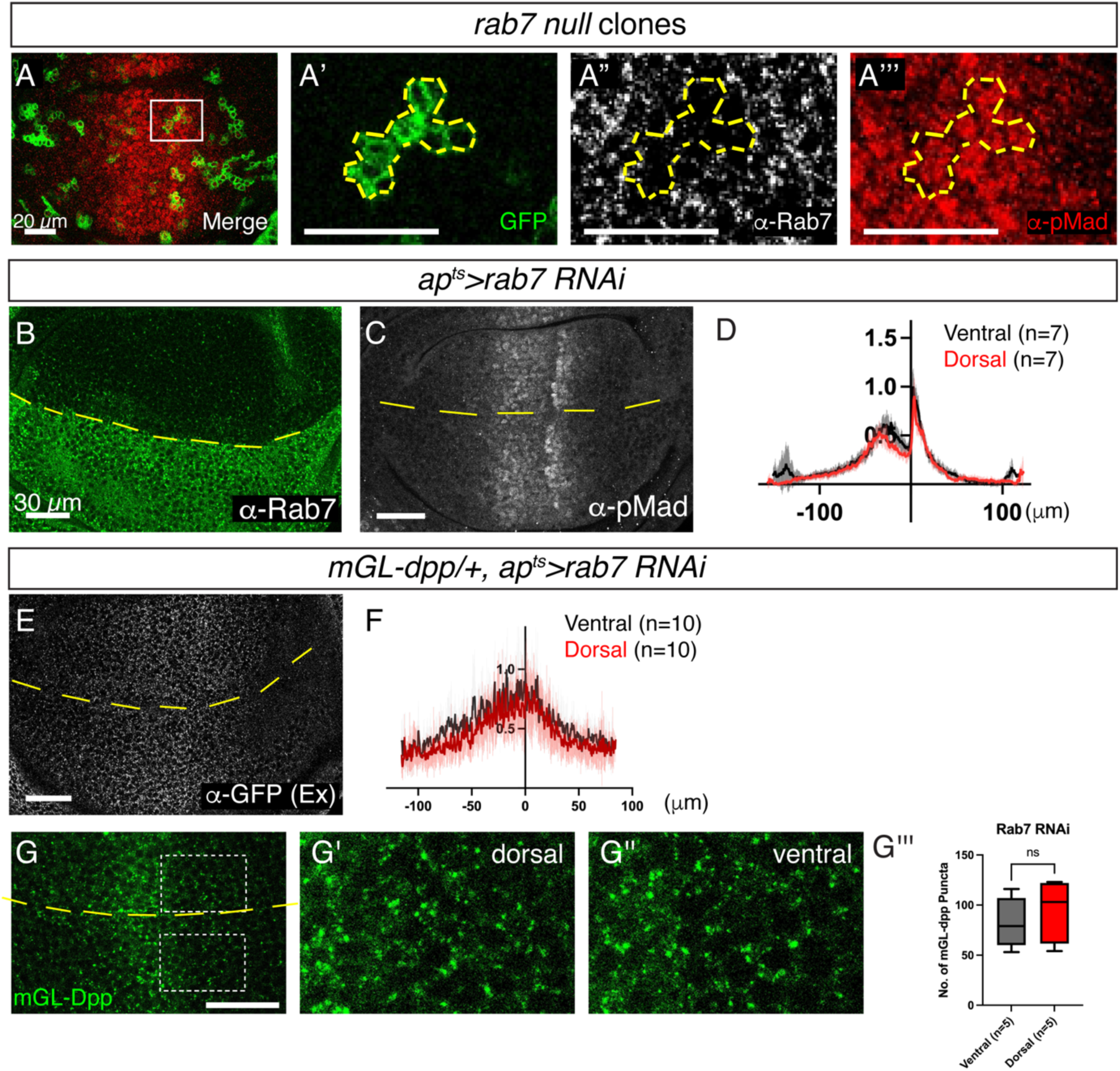
Late endosomal trafficking is not involved in terminating Dpp signaling. (A) Merge (A), GFP signal (A’), α-Rab7 staining (A”), and α-pMad staining (A’’’) of rab7 null clones (labeled by GFP signal) generated by MARCM. (B, C) α-Rab7 staining (B) and α-pMad staining (C) of *ap^ts^>rab7 RNAi* wing disc. (D) Average fluorescence intensity profiles of (C). Data are presented as mean +/-SD. (E) Extracellular a-GFP staining of *mGL-dpp/+, ap^ts^>rab7 RNAi* wing disc. (F) Average fluorescence intensity profiles of (E). Data are presented as mean +/-SD. (G) mGL-Dpp signal from apical side (G), magnified region in the dorsal compartment (G’), and magnified region in the ventral compartment (G”) of *mGL-dpp/+, ap^ts^>rab7 RNAi* wing disc. (G’’’) Comparison of number of mGL-Dpp puncta, ratio-paired t-test with p<0.05 was used for the comparison; non-significant p=0.2083 (n=5).

### Recycling endosome is largely dispensable for extracellular Dpp gradient formation and signaling

mGL-dpp also colocalizes with recycling endosomes (Fig. 2C, D). Since internalized GFP-Dpp revealed by nanobody internalization assay was recycled back to the extracellular space and knocking down Rab4 or Rab11 by RNAi severely affected total GFP-Dpp distribution upon its overexpression ^16^, recycling of Dpp has been proposed to contribute to Dpp gradient scaling. To test if the recycling endosomes are involved in the endogenous Dpp distribution and Dpp signaling, we knocked down Rab4 or Rab11 using the same RNAi lines in the dorsal compartment and investigated Dpp signaling and mGL-dpp distribution. Interestingly, we found that knocking down Rab4 by RNAi did not affect Dpp signaling or extracellular mGL-Dpp distribution except slight decrease in the basal side (Fig. 8A-E). Similarly, knocking down Rab11 by RNAi did not affect Dpp signaling or extracellular mGL-Dpp distribution (Fig.8F-J). Furthermore, while knocking down Rab4 by RNAi did not strongly affect the size and number of intracellular mGL-Dpp puncta (Fig.8K-N), knocking down Rab11 by RNAi caused intracellular accumulation of mGL-Dpp as large puncta (Fig. 8O-R) consistent with defects in endosomal maturation ^40^. These results suggest that Rab4 or Rab11 is not essential for Dpp spreading or signaling.

**Figure 8:**
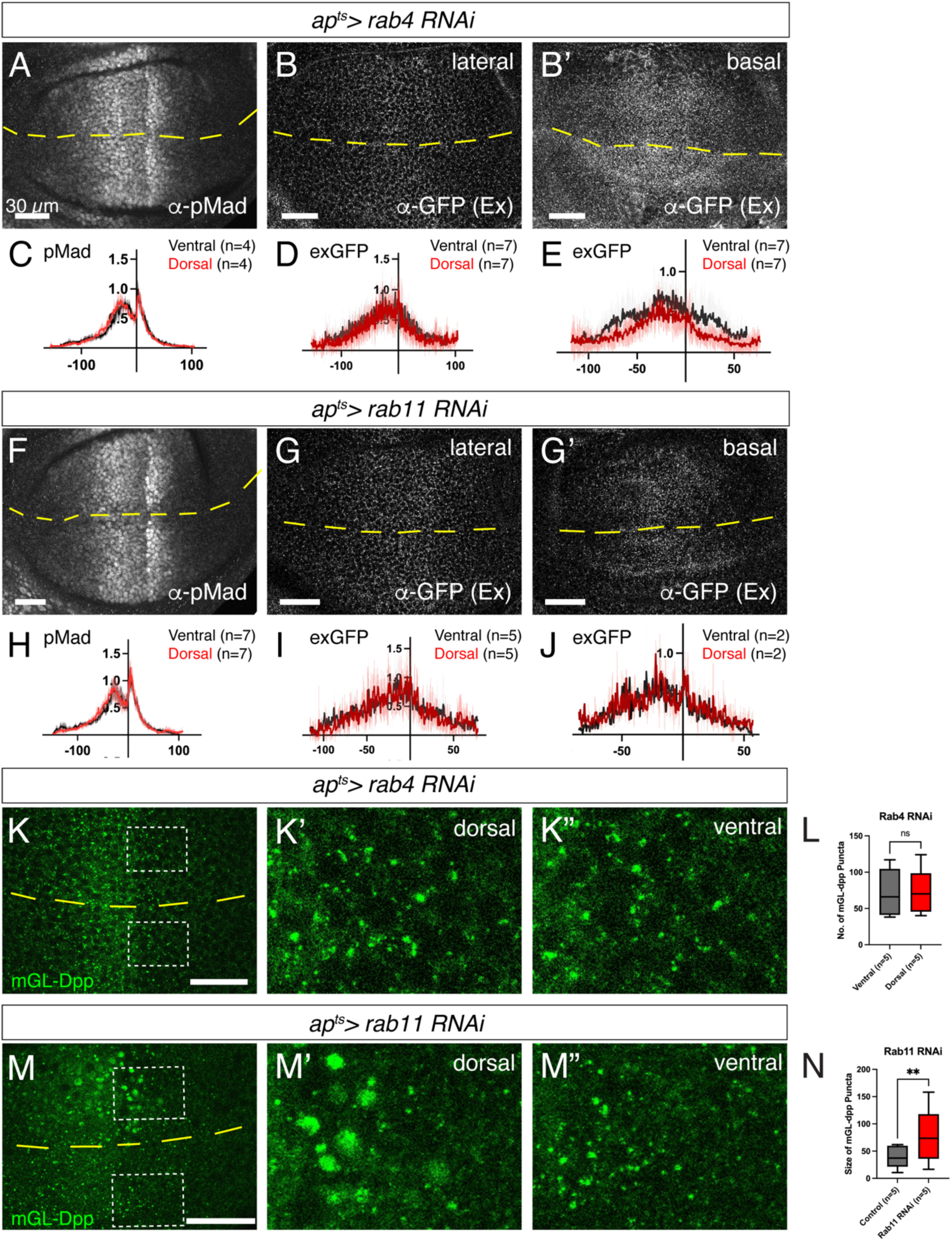
Recycling is largely dispensable for Dpp gradient formation and signaling. (A-B’) α-pMad staining (A), extracellular α-GFP staining in the lateral side (B), and extracellular α-GFP staining in the basal side (B’) of *ap^ts^>rab4 RNAi*. (C-E) Average fluorescence intensity profiles of (A-B’). Data are presented as mean +/-SD. (F-G’) pMad staining (F), extracellular α-GFP staining in the lateral side (G), and extracellular a-GFP staining in the basal side (G’) of *ap^ts^>rab11 RNAi*. (H-J) Average fluorescence intensity profiles of (F-G’). Data are presented as mean +/-SD. (K-L) mGL-Dpp (total) signal of lateral side (K), with magnified region in the dorsal compartment (K’) and ventral compartment (K’’), and comparison between the number of mGL-Dpp puncta in K’ and K’’ (L) in *ap^ts^>rab4 RNAi*. Ratio-paired t-test with p<0.05 was used for the comparison; non-significant p=0.8317 (n=5) (L). (M-N) mGL-Dpp (total) signal of basal side (M), with magnified region in the dorsal compartment (M’) and the ventral compartment (M’’) in *ap^ts^>rab11 RNAi*. (N) Comparison of the size of mGL-Dpp puncta in M’ and M’’. Ratio-paired t-test with p<0.05 was used for the comparison; p=0.006 (n=5).

## Discussion

In this study, we generated novel *dpp* alleles to visualize both extracellular and intracellular Dpp distributions at the physiological condition. Using these alleles, we systematically addressed the role of endocytic trafficking in Dpp distribution and signaling.

Endocytic trafficking has been proposed to regulate extracellular Dpp gradient formation through transcytosis ^15^, recycling ^16^, or a sink ^20–22^. We found that extracellular Dpp distribution expands upon blocking Dynamin or Rab5 (Fig. 4) but is not largely affected upon blocking the following endocytic trafficking such as MVBs (Fig. 6), late endosome (Fig. 7), or recycling endosome (Fig. 8). These results argue against the role of endocytic trafficking in Dpp spreading through transcytosis ^15^ or recycling ^16^ and support the idea that early endocytosis simply acts as a sink for Dpp (Fig. 9). Given that extracellular Dpp distribution expands upon loss of *tkv* ^20,22^, Tkv-mediated internalization of Dpp likely acts as a sink (Fig. 9).

**Figure. 9.**
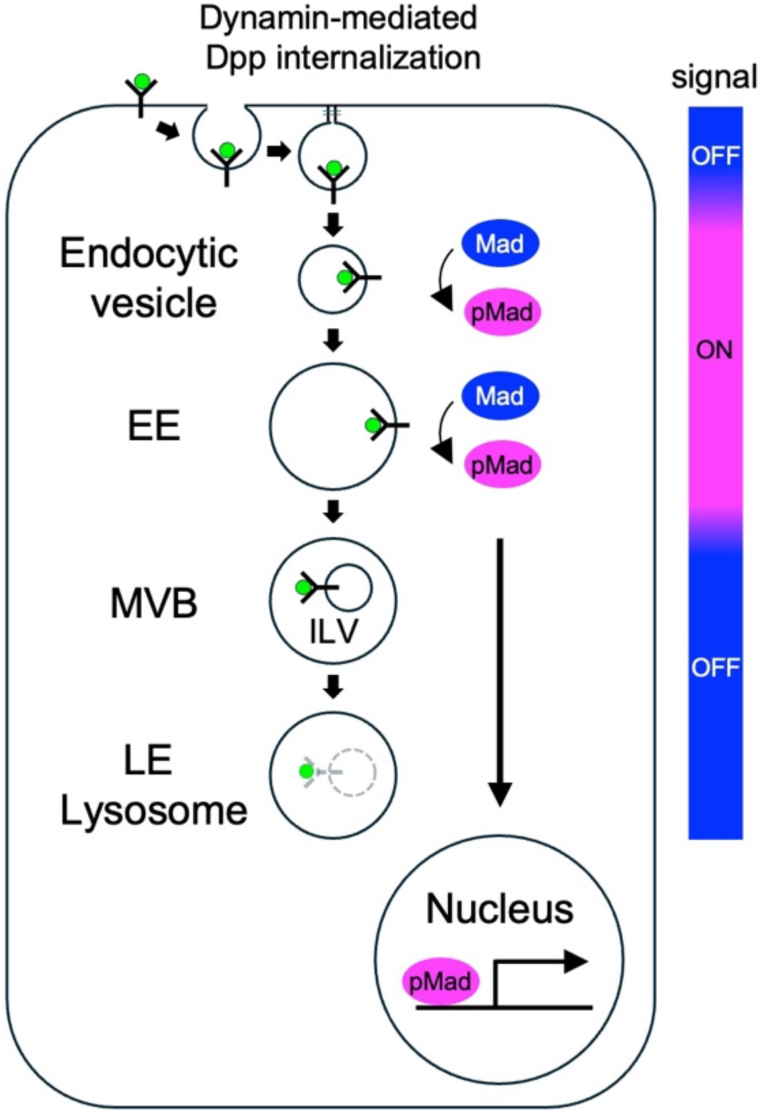
A model for Dpp gradient formation and interpretation by endocytic trafficking. Upon binding to receptor Tkv, extracellular Dpp is internalized by Dynamin-dependent endocytosis. Upon endocytosis, Dpp signaling is turned on in endocytic vesicles before fusing to early endosomes (EE) and turned off through MVB formation, where activated Tkv is physically separated from its target Mad through ILV formation. Duration of Dpp signaling contributes to converting extracellular Dpp gradient into Dpp signaling gradient, since prolonged Dpp signaling by blocking termination of Dpp signaling expands Dpp signaling gradient without affecting extracellular Dpp gradient.

Endocytic trafficking has been also proposed to regulate Dpp signaling. Loss of Dpp signaling by loss of Rab5 indicated that Rab5 is required for Dpp transcytosis and/or Dpp signaling activation^15,31^. However, we found that loss of Rab5 rather caused an increase in Dpp signaling activity due to the impaired downregulation of activated receptors (Fig. 3-5), arguing against these scenarios. Our results indicate that Dpp signaling is activated before fusing to early endosome and is turned off through Rab5-mediated trafficking (Fig. 9). We speculate that the loss of Dpp signaling by loss of Rab5 observed previous studies was in part due to pleiotropic effects. While knocking down Rab5 was performed in a temporally controlled manner using tubGal80ts in this study, Rab5 was constantly knocked down without tubGal80ts in a previous study ^15^. Although the temperature was lowered to express RNAi to reduce pleiotropic effects, we speculate that constant weak expression of dominant negative forms of Rab5 may have nevertheless caused pleiotropic effects affecting Dpp signaling activity. Our results further suggest that Dpp signaling is terminated at the level of MVB formation rather than late endosome-mediated lysosomal degradation of activated receptor (Fig. 6, 7). We speculate that sorting activated receptors into the ILVs itself separates activated Tkv from its target Mad in the cytosol (Fig. 9), contrast to the idea that BMP signal is terminated through lysosomal degradation ^41^. Given that multiple signaling pathways are activated upon blocking MVBs formation, it would be interesting to test if elevated signaling pathways are due to the impaired sorting of the activated receptors into the ILVs or lysosomal degradation. Interestingly, upon blocking MVBs formation, Dpp signaling gradient expanded without affecting extracellular Dpp gradient (Fig. 6F-H), compromising the interpretation of extracellular Dpp gradient. Thus, duration of Dpp signaling plays a critical role in converting extracellular Dpp gradient to the proper Dpp signaling activity gradient (Fig. 9).

### Novel *dpp* alleles to visualize endogenous Dpp morphogen gradient

Dpp morphogen gradient has been intensively studied using GFP-Dpp. When expressed in the anterior stripe of cells, the main *dpp* source, GFP-Dpp showed highest fluorescent signal in the source cells and shallow graded signal in both sides as punctate signal ^15,24^. While the punctate signal was shown to be mainly from the internalized GFP-Dpp ^23–25^, extracellular staining revealed distinct extracellular-specific Dpp morphogen gradient ^22^. Using FRAP and FCS, the kinetics of Dpp gradient formation have been measured, including diffusion coefficient, degradation rates, and decay length ^23,25^. However, given the unphysiological level of overexpression (estimated 400 times higher than the physiological level) ^16^, it has been questioned how much the obtained results from overexpression of Dpp reflects the mechanisms underlying the endogenous Dpp gradient formation ^20^. Indeed, contrast to the results obtained using overexpression of GFP-Dpp ^16^, we could not detect severe defects in extracellular Dpp distribution or Dpp signaling upon loss of Rab4 or Rab11 (Fig. 8). Nevertheless, these results suggested that Dpp gradient consists of extracellular (bound and unbound on the cell surface) and internalized populations.

Recently, with the advances in genome engineering methods, it has become possible to insert a tag in *dpp* locus ^7,16,42^. Endogenous *GFP-dpp* allele revealed that the fluorescent signal was too low to visualize the graded Dpp distribution (Fig. 1A) and to apply FRAP assay to measure the parameters of Dpp gradient formation ^16^. Similarly, an endogenous *HA-dpp* allele revealed a shallow extracellular HA-Dpp distribution and the conventional immunostaining failed to visualize the Dpp distribution outside the main source cells ^7^. The nanobody internalization assay was able to visualize the internalized Dpp but it is not clear if the nanobody bound GFP-Dpp reflects the functional ligand that undergoes proper endocytic trafficking.*mGL-dpp* and *mSC-dpp* alleles can overcome these shortcomings. These alleles are functional at least during wing development and their brighter fluorescent signal allows for visualization of the endogenous Dpp distribution (mostly internalized Dpp) without any manipulation (Fig. 1). Using the *mGL-dpp* allele also allows for visualization of the extracellular Dpp distribution through anti-GFP antibody staining. FRAP assays, morphotrap, and live imaging have already been successfully applied to characterize the role of mGL-Dpp in Drosophila ^43^. By applying these assays in the wing disc, it would be of interest to re-investigate the kinetics of Dpp morphogen gradient formation under physiological conditions.

### Materials and methods Fly stocks

Flies for experiments were kept in standard fly vials containing polenta and yeast. Embryos from fly crosses for experiments including Gal80ts were collected for 24h and kept at 18°C, until shifted to 29°C prior to dissection of 3^rd^ instar larvae. To induce *Rab5*^2^ clones, third instar larvae were subjected to heat shock (37°C) for 8 minute and incubated at 25°C for 24 hours prior to dissection. The following fly lines were used: *shibire^ts^*^1^ (BDSC 7068), *mGL-dpp* (this study), *mSC-dpp* (this study), *ap-Gal4* (M. Affolter), *tub-GAL80TS* (M. Affolter), *tkv-3xHA* (G. Pyrowolakis), *tkv-YFP* (G. Pyrowolakis), *tkv-1xHAeGFP* (G. Pyrowolakis), *brk^XA^* (G. Campbell & A. Tomlinson), *UAS-rab5-RNAi* (BDSC 30518, VDRC 34096, 103945), *UAS-rab5.S43N* (BDSC 42703 & 42704), *UAS-rab4 RNAi* (VDRC 24672), *UAS-rab11-RNAi* (VDRC 22198), *UAS-vps4-RNAi* (VDRC 105977), *UAS-tsg101-RNAi* (BDSC 35710), *UAS-shrub-RNAi* (BDSC 38305*), UAS-rab7-RNAi* (BDSC 27051), *dpp-LacZ* (M.Affolter), *UAS-LOT-deGradHA* (G. Pyrowolakis & M. Affolter), *rab5-eYFP* (BDSC 62543*), rab7-eYFP* (BDSC 62545), *rab4-eYFP* (BDSC 62542*), rab11-eYFP* (BDSC 62549), *FRT82b, rab7^Gal^*^4^*^-Knock-in^*null allele (P. R. Hiesinger), *hsFlp,UAS-GFP,w;FRT42D,tub-Gal80;tub-Gal4,FRT82B,tub-Gal80* (BDSC 86318), *hsFlp;tub>CD2>Gal4,UAS-lacZ* (B. Bello), *hsFlp, rab5*^2^*, FRT40* (BDSC 42702), *yw, dpp^d^*^8^ and *dpp^d^*^12^ are described from Flybase.

### Genotypes by figures

|  |
| --- |
| Fig.1A; *GFPdpp/Cyo, p23* |
| Fig.1B; *mGL-dpp/+* |
| Fig.1C; *mSC-dpp/+* |
| Fig.1D; *JAX; mGL-dpp/mGL-dpp* |
| Fig.1E; *JAX; mSC-dpp/mSC-dpp* |
| Fig.1F; *HA-dpp/HA-dpp* |
| Fig.1G; *JAX; HA-dpp/HA-dpp* |
| Fig.1H; *JAX; mGL-dpp/mGL-dpp* |
| Fig.1I; *JAX; mSC-dpp/mSC-dpp* |
| Fig.1K; *mGL-dpp/+* |
| Fig.2A; *mSC-dpp / rab5-eYFP* |
| Fig.2B; *mSC-dpp / +; rab7-eYFP / +* |
| Fig.2C; *mSC-dpp / rab4-eYFP* |
| Fig.2D; *mSC-dpp / +; rab11-eYFP / +* |
| Fig. 3A; *shi*<sup>TS</sup>/+ (2h heat shock at 34°C) |
| Fig. 3B; *shi*<sup>TS</sup> (2h heat shock at 34°C) |
| Fig.3D, E; *HA-dpp, ap-Gal4 / +; UAS-rab5-RNAi* (30518) / *tub-Gal80ts* (29h at 29°C) |
| Fig.3G; *HA-dpp, ap-Gal4 / +; UAS-rab5-RNAi* (34096) / *tub-Gal80ts* (29h at 29°C) |
| Fig.3H; *HA-dpp, ap-Gal4 / +; UAS-rab5-RNAi* (103945) / *tub-Gal80ts* (24h at 29°C) |
| Fig.3I; *HA-dpp, ap-Gal4 / UAS-rab5.S43N* (42703); *tub-Gal80ts / +* (18h at 29°C) |
| Fig.3J; *Ollas-dpp, ap-Gal4 / +; UAS-rab5.S43N* (42704)/ *tub-Gal80ts* (13.5h at 29°C) |
| Fig.3K; *hsFlp, rab5*<sup>2</sup> *FRT40/arm-LacZ, m(2)Z FRT40* |
| Fig.4A; *brk*<sup>XA</sup>; *dpp*<sup>d8</sup>, *ap-Gal4 / dpp*<sup>d12</sup>; *UAS-rab5-RNAi* (30518) / *tub-Gal80ts* (29h at 29°C) |
| Fig.4B; *apGal4 / dpp-LacZ; UAS-rab5-RNAi* (30518) / *tub-Gal80ts* (29h at 29°C) |
| Fig.4C,D; *mGL-dpp*, *apGal4* / +; *UAS-rab5-RNAi* (30518) / *tub-Gal80ts* (29h at 29°C) |
| Fig.4F; *shi<sup>TS</sup>* / +; *mGL-dpp* / + (2h heat shock at 34°C) |
| Fig.4G; *shi<sup>TS</sup>*; *mGL-dpp* / + (2h heat shock at 34°C) |
| Fig.4I,J; *dpp-mGL*, *apGal4* / +; *UAS-rab5-RNAi* (30518) / *tub-Gal80ts* (29h at 29°C) |
| Fig.4K, L; *tkv-YFP*, *apGal4*; *UAS-rab5-RNAi* (30518) / *tub-Gal80ts* (29h at 29°C) |
| Fig.5A; *yw*, *tkv-HA-eGFP*, *apGal4* / +; *tub-Gal80ts* / + (29h at 29°C) |
| Fig.5B; *tkv-HA-eGFP*, *apGal4* / +; *UAS-rab5-RNAi* (30518) / *tub-Gal80ts* (29h at 29°C) |
| Fig.5C; *tkv-HA-eGFP*, *apGal4* / +; *UAS-rab5-RNAi* (30518), *tub-Gal80ts* / *UAS-deGradHA* (29h at 29°C) |
| Fig.5D; *tkv-HA-eGFP*, *apGal4* / +; *UAS-deGradHA* / *tub-Gal80ts* (29h at 29°C) |
| Fig.6A; *tkv-YFP*, *apGal4*; *UAS-tsg101-RNAi* (35710) / *tub-Gal80ts* (44h at 29°C) |
| Fig.6B; *tkv-YFP*, *apGal4* / *UAS-shrub-RNAi* (38305); *tub-Gal80ts* / + (28h at 29°C) |
| Fig.6C, D; *tkv-YFP*, *apGal4* / *UAS-vps4-RNAi* (105977); *tub-Gal80ts* / + (30h at 29°C) |
| Fig.6E; *tkv-YFP*, *apGal4*; *UAS-rab7-RNAi* (27051) / *tub-Gal80ts* (42h at 29°C) |
| Fig.6F, H; *mGL-dpp*, *apGal4* / *UAS-vps4-RNAi* (105977); *tub-Gal80ts* / + (30h at 29°C) |
| Fig.7A; *hsFlp*, *UAS-GFP*; *FRT82b*, *tub-Gal4* / *FRT82b rab7 Gal4-Knock-In* |
| Fig.7B, C; *HA-dpp*, *apGal4*; *UAS-rab7-RNAi* (27051) / *tub-Gal80ts* (42h at 29°C) |
| Fig.7C, G; *mGL-dpp*, *apGal4*; *UAS-rab7-RNAi* (27051) / *tub-Gal80ts* (42h at 29°C) |
| Fig.8A, B; *HA-dpp*, *ap-Gal4* / +; *UAS-rab4-RNAi* (24672) / *tub-Gal80ts* (42h at 29°C) |
| Fig.8F, G; *HA-dpp*, *ap-Gal4* / +; *UAS-rab11-RNAi* (22198) / *tub-Gal80ts* (42h at 29°C) |
| Fig.8K; *HA-dpp*, *ap-Gal4* / +; *UAS-rab4-RNAi* (24672) / *tub-Gal80ts* (42h at 29°C) |
| Fig.8M; *HA-dpp*, *ap-Gal4* / +; *UAS-rab11-RNAi* (22198) / *tub-Gal80ts* (42h at 29°C) |

### Generation of *mGL-dpp* and *mSC-dpp*

The detail procedure to generate endogenously tagged *dpp* alleles were previously reported^7^. In brief, utilizing the attP sites in a MiMIC transposon inserted in the dpp locus (MiMIC dppMI03752, BDSC 36399), about 4.4 kb of the dpp genomic sequences containing the second (last) coding exon of dpp including a tag and its flanking sequences was inserted in the intron between dpp’s two coding exons. The endogenous exon was then removed using FLP-FRT to keep only the tagged exon. mGL (mGreenLantern ^27^) was inserted after the last processing site to tag all the Dpp mature ligands. mGL-dpp homozygous flies show no obvious phenotypes.

## Immunohistochemistry

### Visualization of mGL-Dpp and mSC-Dpp

To visualize the (total) mGL-Dpp and mSC-Dpp signal, third instar larvae were dissected in ice-cold Phosphate Buffered Saline (PBS). The dissected larvae were washed with HCl with pH 3.0 following the acid wash protocol ^16^ to remove the extracellular proteins prior to fixation in 4.0% Paraformaldehyde (PFA) for 25min on a shaker at room temperature (25°C). The discs were washed three times for ten minutes with PBS at 4°C and mounted in Vectashield on glass slides.

### Total staining

Third instar larvae were dissected in ice-cold Phosphate Buffered Saline (PBS) and fixed in 4.0% Paraformaldehyde (PFA) for 25min on a shaker at room temperature (25°C). After fixation, the discs were washed three times for ten minutes with PBS at 4°C, and three times with PBST (0.3% Triton-X in PBS) to permeabilize the tissues. The discs were then blocked in 5% normal goat serum (NGS) in PBST for 30min. The primary antibodies were added to 5% NGS in PBST for incubation of the discs at 4°C overnight. The next day, the primary antibody was carefully removed, and the samples were washed three times with PBST. Secondary antibodies were added to5 % NGS in PBST and the discs were incubated for 2h in the dark at room temperature. At last, the samples were washed three times for 15 minutes with PBST at room temperature, two times with PBS, and mounted in Vectashield on glass slides.

### Extracellular staining

Wing discs from third instar larvae were dissected in ice-cold Schneider’s Drosophila medium (S2). The discs were then blocked in cold 5% NGS in S2 medium on ice for 10min. The blocking solution was carefully removed, and the primary antibody was added for 1h on ice. To ensure an even distribution of the primary antibody, the tubes were tapped every 10min during the incubation time. The antibody was then removed, and the samples were washed at least 6 times with cold S2 medium and another two times with cold PBS to remove excess primary antibody. Wing discs were then fixed with 4% PFA in PBS for 25min on the shaker at room temperature (25°C). After fixation the protocol continued as described in total staining.

### Acid wash

The protocol was adapted from ^16^. To remove the extracellular proteins prior to fixation, the dissected wing discs were washed three times ten seconds with ice-cold Schneider’s Drosophila medium (S2), pH dropped down to 3 by HCl. To remove the stripped membrane-bound proteins, the discs were washed three times 15min with ice-cold S2 medium (pH 7.4) and fixed in 4% PFA.

### Antibodies

Primary antibodies: Rabbit anti-phospho-Smad 1/5 (Cell signaling 9516S; 1:200), mouse anti-patched (DSHB; 1:40), mouse anti-wingless (4D4, DSHB; 1:120), rabbit anti-GFP (Abcam ab6556; 1:2000 for total staining, 1:200 for extracellular staining,), guinea pig anti-rab5 (provided by Akira Nakamura; 1:1000), rabbit anti-rab11 (provided by Akira Nakamura; 1:8000), mouse anti-rab7 (DSHB; 1:30), mouse anti-ubiquitin (Enzo PW8810-0100; 1:1000), mouse anti-beta galactosidase (Promega Z378825580610; 1:500), guinea pig anti-brk (provided by from Gines Morata; 1:1000), mouse anti-V5 (Invitrogen; 1:5000).

The following secondary antibodies were used at 1:500 dilutions in this study: Goat anti-rabbit IgG (H+L) Alexa Fluor^TM^ 488 (A11008 Thermo Fischer), goat-anti-rabbit IgG (H+L) Alexa Fluor^TM^ 568 (A11011 Thermo Fischer), goat-anti-rabbit IgG (H+L) Alexa Fluor^TM^ 680 (A21109 Thermo Fischer), goat anti-mouse IgG (H+L) Alexa Fluor^TM^ 488 (A11001 Thermo Fischer), goat anti-mouse IgG (H+L) Alexa Fluor^TM^ 568 (A11004 Thermo Fischer), goat anti-mouse IgG (H+L) Alexa Fluor^TM^ 680 (A10038 Thermo Fischer), goat-anti-guinea pig IgG (H+L) Alexa Fluor^TM^ 568 (A11075 Thermo Fischer), goat-anti-guinea pig IgG (H+L) DyLight 680 (SA5-10098 Invitrogen).

### Imaging

Wing imaginal discs were imaged using a Leica SP5-II MATRIX and an Olympus Spinning Disk (CSU-W1), and images were analyzed using Fiji (ImageJ). Figures were obtained using OMERO and Adobe Illustrator.

### Quantification of pMad and extracellular mGL-dpp intensity

To quantify the intensity of pMad and extracellular mGL-dpp gradient in the images, an average intensity of three sequential stacks was created using Fiji ImageJ (v1.53c). Each signal intensity profile collected in Excel (Ver. 16.51) was aligned along A/P compartment boundary (based on anti-Ptc or pMad staining) and average signal intensity profile from different samples was generated and plotted by the script (wing_disc-alignment.py). The average intensity of the samples and the control were then compared using the script (wingdisc_comparison.py). Both scripts were generated by E. Schmelzer and can be found on: https://etiennees.github.io/Wing_disc-alignment/. The resulting signal intensity profiles (mean with SD) were generated on GraphPad Prism software (v.9.3.1(471)). Figures were prepared using OMERO (ver5.9.1) and Adobe Illustrator (24.1.3).

### Quantification of mGL-dpp and Tkv-YFP positive puncta

To measure the number particles an average intensity of 3 z-stacks from the images were created using Fiji ImageJ. The total area of controls and samples in which the particles were counted had a width of 20.16 and height of 34.17 microns. The number and area of the particles were measured by the built-in “Analyze Particles” plug-in in Fiji. The data were used to make the graphs on GraphPad Prism. A ratio-paired t-test (p<0.05) was used for statistical analysis.

### Reproducibility

All experiments were independently repeated at least two time, with consistent results. Statistical significance was assessed by the GraphPad Prism software (v.9.3.1(471)).

## Acknowledgements

The authors would like to thank Markus Affolter for his continuous support throughout the course of this project. We thank Developmental Studies Hybridoma Bank (DSHB) at The University of Iowa for providing us with the primary antibodies, and Bloomington Drosophila Stock Center (BDSC) for providing us with fly stocks. We would also like to thank Dr. Giorgos Pyrowolakis, Prof. Peter Robin Hiesinger and Prof. Isabel Guerrero for providing us with fly lines and Prof. Akira Nakamura for providing us with primary antibodies. We thank Dr. Etienne Schmelzer for providing us with scripts for quantifications. We would like to thank Bernadette Bruno, Gina Evora, Karin Mauro and Dario Dörig for their constant and reliable supply of the world’s best fly food. We thank the Biozentrum Imaging Core Facility (IMCF), especially Dr. Oliver Biehlmaier, Dr. Alexia Loyton-Ferrand, Dr. Sara Roig, Dr. Kai Schleicher, Laurent Guerard, Nikolaus Ehrenfeuchter and Dr. Sébastien Herbert for their constant support with the microscopes and image analysis.

---

**Figure S1:**
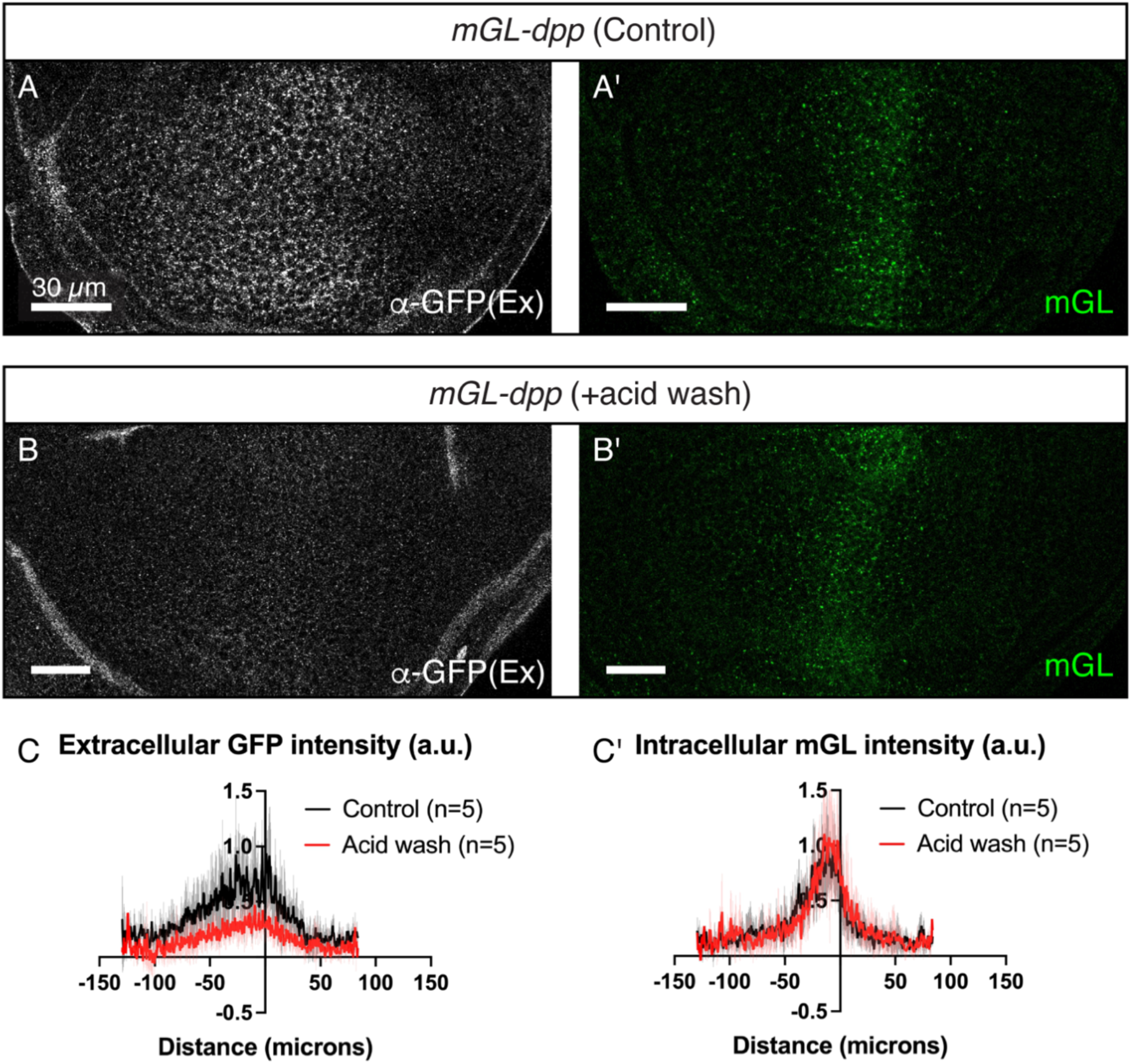
Acid wash removes extracellular mGL-Dpp but does not alter the intracellular mGL-Dpp signal. (A) Extracellular mGL-Dpp observed through an α-GFP antibody staining in the control condition in absence of the acid wash, (A’) total mGL-Dpp signal in absence of the acid wash, (B) Extracellular mGL-Dpp observed through an α-GFP antibody staining after the acid wash, (B’) Total mGL-Dpp signal after the acid wash. (C-C’) Quantification of the extracellular mGL-Dpp signal intensity in A and B (C), and the intracellular mGL-Dpp signal in A’ and B’ (C’).

**Figure. S2:**
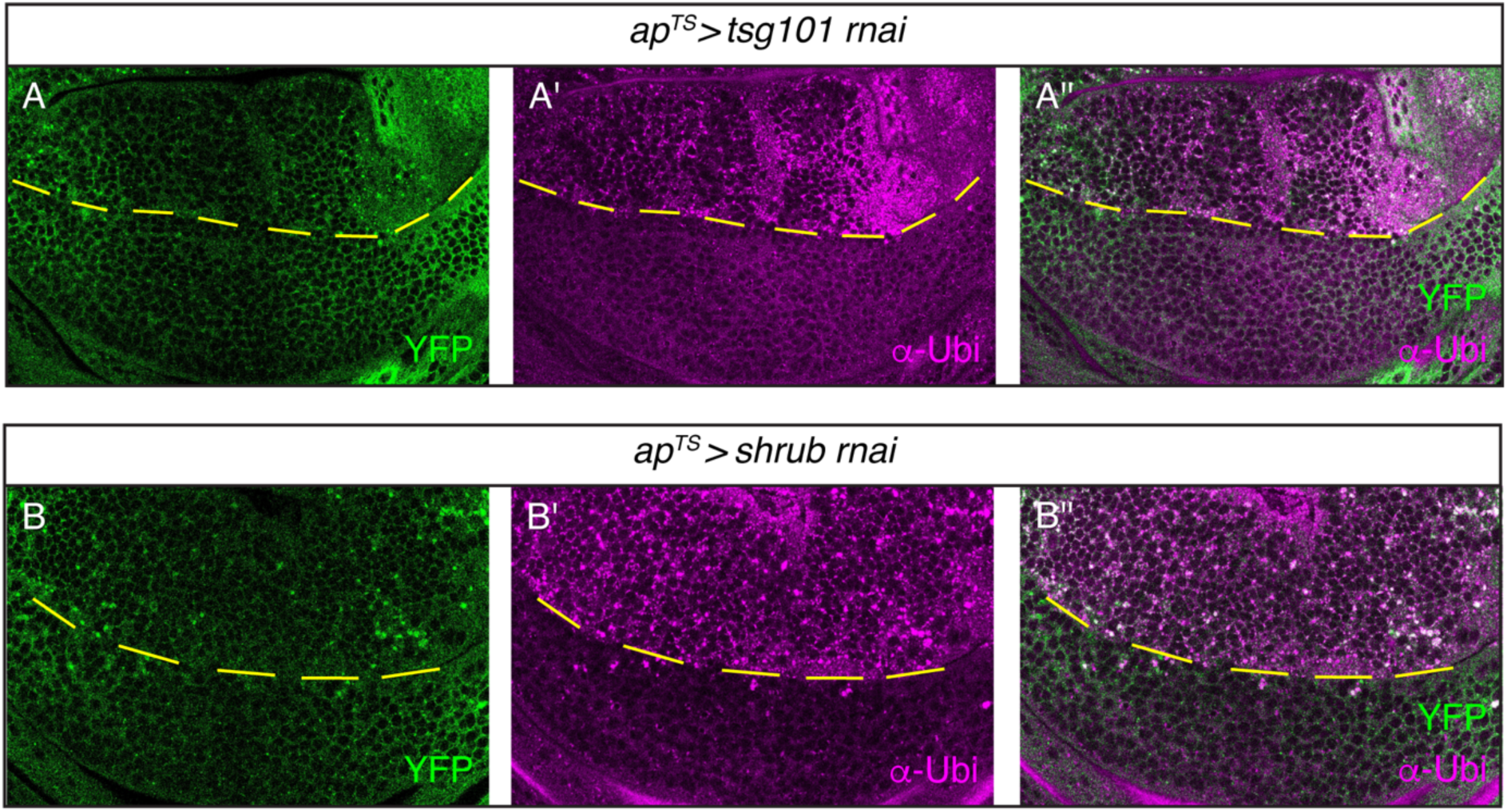
Knocking down the ESCRT components TSG101 and shrub leads to an accumulation of Tkv and ubiquitin in puncta. (A-A’’) Tkv-YFP signal (A), ubiquitin signal (A’) and the merged image (A’’) in *ap^TS^>tsg101 rnai* flies. (B-B’’) Tkv-YFP signal (B), ubiquitin signal (B’) and the merged image (B’’) in *ap^TS^>shrub rnai* flies.

